# Transertion is used for localized expression and assembly of *Vibrio parahaemolyticus* T3SS2

**DOI:** 10.1101/2022.02.23.481666

**Authors:** Karan Gautam Kaval, Suneeta Chimalapati, Sara Siegel, Nalleli Garcia Rodriguez, Ankur B. Dalia, Kim Orth

## Abstract

Using environmental cues, bacteria commit to the assembly of transmembrane complexes such as the type III secretion system 2 (T3SS2), a membrane-bound, syringe-like secretory apparatus used during infection to inject host cells with virulence factors. Here we report *Vibrio parahaemolyticus* uses transertion, localized transcription, translation, and membrane insertion, to assemble its T3SS2. Upon binding bile acids, the membrane bound receptor and transcription factor VtrA/VtrC captures the T3SS2 pathogenicity island at the inner membrane. Activated VtrA/VtrC induces production of VtrB, the membrane bound master T3SS2 transcriptional regulator. VtrB then induces the membrane-proximal T3SS2 genes to undergo transertion for assembly of the membrane inserted secretion machinery. Transertion is a process that can be used for the efficient assembly of membrane-bound molecular complexes in response to extracellular signals.

**One-Sentence Summary:** Localized transcription, translation, and membrane insertion of multi-protein complexes in bacteria in response to host cues.

## Main Text

In Gram-negative bacteria, one-component regulators unlike those of two-component systems are predominantly cytoplasmic. However, some of these transcription factors are bitopic, characterized by the presence of a ligand-binding periplasmic sensor domain, and a cytoplasmic helix-turn-helix DNA binding domain connected via a single-pass transmembrane region (*1*). Previous studies have shown that once these receptors are activated by their respective ligands, they use a diffusion/capture mechanism to localize their genomic target promoters to the membrane, allowing them to bind to and initiate transcription (*2, 3*). The Gram-negative gastrointestinal pathogen, *Vibrio parahaemolyticus*, employs such a membrane bound heterodimeric one-component system, VtrA/VtrC, to activate expression of the genes encoding the syringe-like Type III Secretion System 2 (T3SS2). Upon induction with bile acids, VtrA/VtrC mediates the injection of specialized effector proteins into host cells to overcome their cellular processes (*4-6*). Herein, we report that the membrane assembly of the T3SS2 apparatus in *V. parahaemolyticus* is driven by the long theorized transertion mechanism, a term for the concurrent transcription, translation, and membrane insertion of biosynthetic products initiated by membrane localization of their encoding genes (*7-10*). Assembly of the *V. parahaemolyticus* T3SS2 by transertion is induced in two steps, by two membrane tethered transcription factors that work in tandem. Binding of bile acids to the periplasmic domain of the VtrA/VtrC complex induces dimerization and activation of its cytoplasmic DNA-binding helix-turn-helix domain (HTH), allowing it to capture the T3SS2 pathogenicity island at the membrane via the encompassed *vtrB* promoter (fig. S1A). This membrane capture initiates the first transertion step allowing for the expression, assembly and activation of the VtrB transcription factor at the membrane (fig. S1B). As a master regulator of the T3SS2, VtrB then carries out the second transertion step by binding to its target promoters within the membrane associated T3SS2 pathogenicity island and inducing localized production of the T3SS2 components at the membrane. These two transertion steps enable efficient assembly of the T3SS2 apparatus with its chaperone-bound effectors (fig. S1C, Fig 2A).

To better understand the effect of bile acid-mediated VtrA/VtrC activation on the membrane proximity of the *vtrB* genomic locus, a locus-tagging and tracking methodology based on a minimal bacterial plasmid-partitioning Par system was employed (*11*). In the *V. parahaemolyticus* CAB2 strain background ((*12*); Table S1), we introduced a *parSMT1* sequence from the *E. coli* MT1 plasmid in the intergenic region downstream of the *vtrB* genomic locus. The cognate fluorescently labelled ParBMT1 was expressed under the control of an arabinose-inducible promoter, to tag and track the genomic *vtrB* locus (Fig. 1A). These cells were cultured in the presence or absence of the activating bile salt, taurodeoxycholate (TDC). Localization of the fluorescently labeled *vtrB* loci within the cells (Fig. 1B) was observed and quantified by measuring their distances from the membrane (Fig. 1C and D). Distribution of normalized distances of the *vtrB* loci from the membrane (p’) revealed membrane bias of this loci in cells grown with TDC as compared to those in cells grown in noninducing conditions (Fig. 1E). A neutral locus (*0*.*458Mb* on chromosome 2) located outside the T3SS2 pathogenicity island was monitored simultaneously using an analogous Par system from the *E. coli* P1 plasmid (*11*) (fig. S2A and B). The neutral locus displayed no differences in its distribution within cells cultured either with or without TDC (fig. S2C right panel). Collectively, these data demonstrate that upon bile-acid induction, activated VtrA/VtrC specifically captures the *vtrB* locus at the inner membrane.

**Figure 1.**
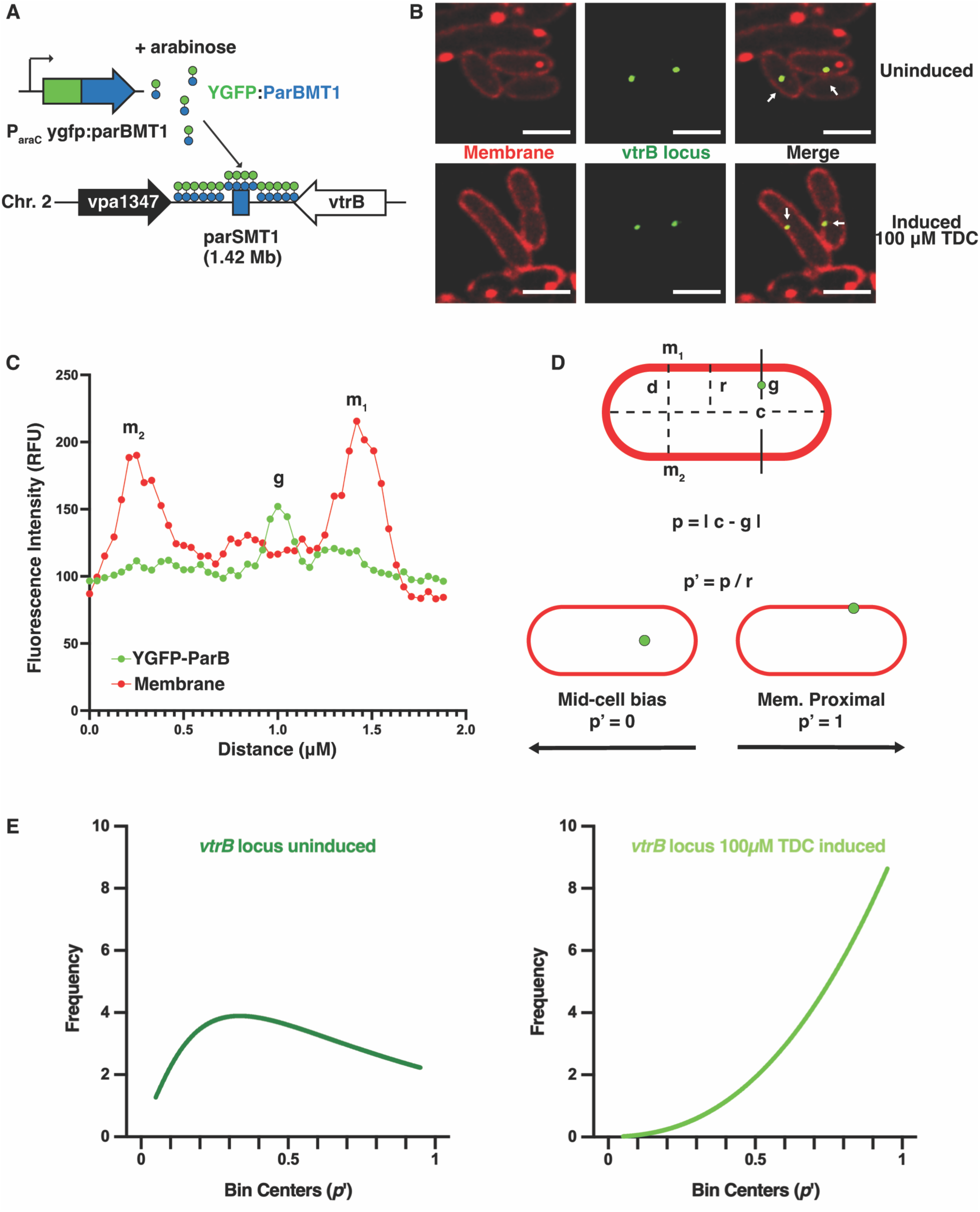
Bile induction of VtrA/VtrC leads to inner membrane capture of the *vtrB* locus. **A**. Illustration depicting *vtrB* genomic locus-tagging using the *parsMT1-*ParBMT1 system in *V. parahemolyticus*. **B**. Confocal micrographs showing the intracellular localization of the *vtrB* genomic loci (green, pointed to by white arrows) in relation to the membrane (red) in *V. parahaemolyticus* SS2B5 cells cultured in non-inducing or inducing (100 µM TDC) conditions with 0.02% arabinose supplementation for YGFP-ParBMT1 expression. Membranes were stained with nile red. Scale bar = 2 µm. **C**. Fluorescence intensity profile of a *V. parahaemolyticus* SS2B5 cell grown in non-inducing conditions for *vtrB* locus membrane proximity quantification. Intensity peaks “m_1_” and “m_2_” denote lateral membrane fluorescence intensities, while “g” corresponds to the intensity of the YGFP-ParBMT1 punctum (*vtrB* locus). **D**. Illustration describing methodology for the normalization of *vtrB* locus membrane proximity (p’) where g = *vtrB* locus (µm), m_1_ = lateral wall 1 (µm), m_2_ = lateral wall 2 (µm), were calculated using the fluorescence intensity profile for each YGFP-ParBMT1 punctum. For subsequent calculations, d = diameter (µm), c = cell center (µm), and r = cell radius (µm). **E**. Frequency distribution plots with nonlinear regression (lognormal) analyses of normalized *vtrB* loci distances (p’) in *V. parahaemolyticus* SS2B5 cells, cultured in non-inducing (left panel) and inducing (100 µM TDC, right panel) conditions. N=30 *vtrB* loci per condition.

Like VtrA/VtrC, VtrB is an inner membrane protein with a cytoplasmic N-terminal HTH DNA-binding domain (*13*), and is referred to as a master regulator for the T3SS2, that binds the promoters of genes encoding T3SS2 components. Therefore, to visualize VtrB-mediated transertion of the T3SS2, VtrB was expressed from its native locus with its periplasmic component tagged with a monomeric superfolding GFP that can efficiently fold and fluoresce within the periplasmic oxidative niche (*14*). The membrane proximity of the *vtrB* genomic locus was simultaneously visualized in this background using the *parS*-ParBMT1 locus-tagging system described previously, except with the YGFP tag of ParBMT1 swapped for CFP. Colocalization of VtrB (false-colored blue) with the *vtrB* locus (false-colored green) at the inner membrane was observed in cells of this strain induced with TDC (Fig. 2B). However, in the absence of inducing conditions, the *vtrB* locus displayed mid-cell bias with absence of any observable VtrB expression (Fig. 2B). These observations indicate that TDC-induction of the VtrA/VtrC transcription factor results in the localized expression and assembly of VtrB at the inner membrane via transertion.

**Figure 2.**
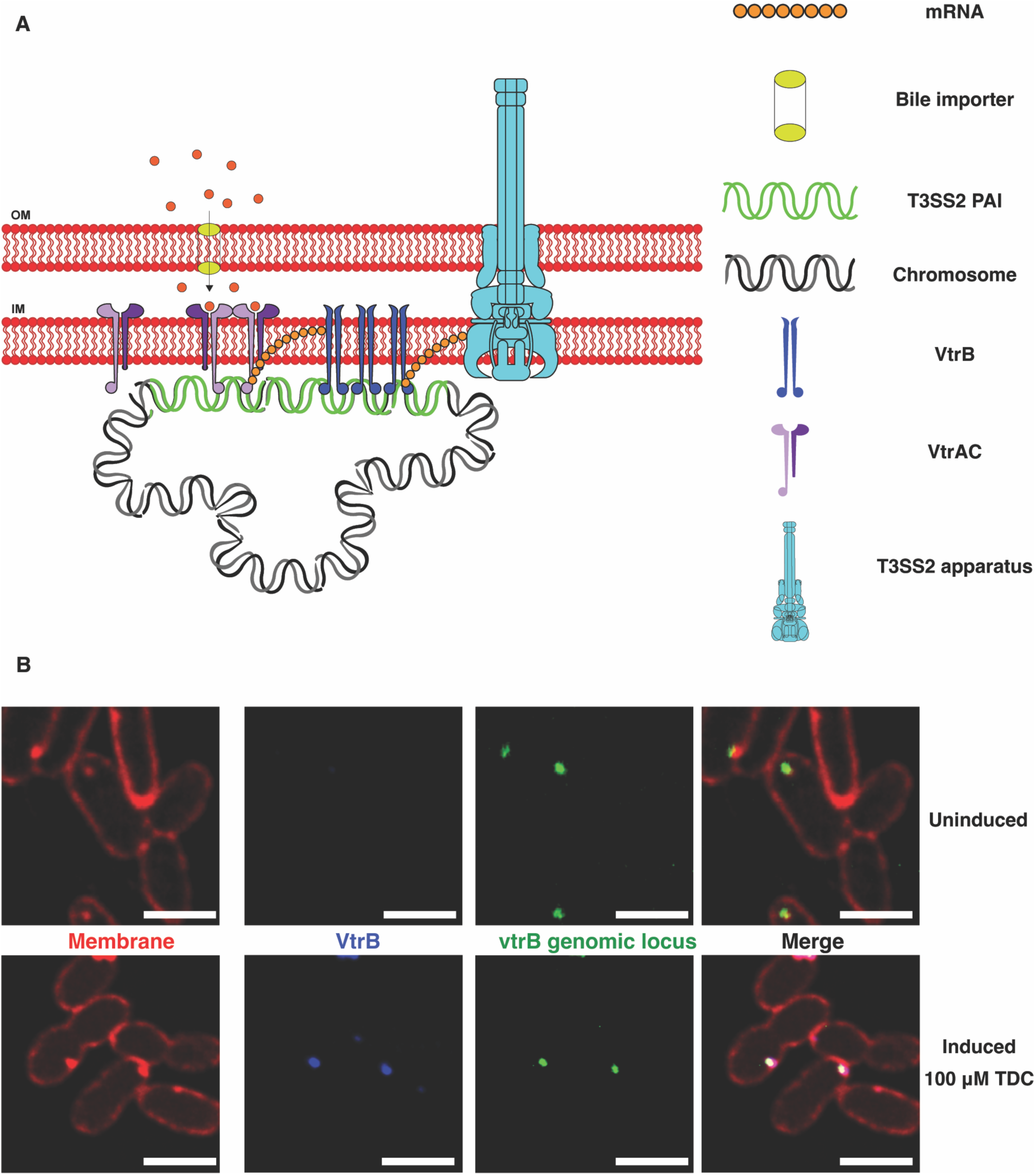
VtrB is assembled at the site of *vtrB* locus capture in the inner membrane. **A**. Illustration depicting the assembly of the T3SS apparatus at the membrane, driven by the hypothesized transertion mechanism upon bile acid-induction of the VtrA/VtrC membrane-bound transcription factor **B**. Confocal micrographs showing expression and colocalization of VtrB (false-colored blue) with the *vtrB* genomic locus (false-colored green) at the membrane in *V. parahaemolyticus* SC295 cells cultured in inducing (100 µM TDC) vs. non-inducing conditions. Scale bar = 2 µm.

The T3SS2 injectosome is a multiprotein complex consisting of ∼19 core structural components, all of which are essential for the functionality of this apparatus (*15, 16*). Of these components, the needle protein is the one that links the membranes of the bacteria and the host cell by forming a filament through the extracellular space, allowing for insertion of the injection apparatus into the host membrane to facilitate transfer of bacterial effectors into the host cytosol (*17*). While most of the T3SS2 structural components have been characterized, the identity of the needle filament protein, in addition to four others, remains obscure due to low sequence similarity with other T3SS needle proteins (*15, 16*). Using sensitive sequence similarity detection methods (HHPRED), we found the T3SS2 protein, Vpa1343, to confidently identify the secretion system needle proteins from *Yersinia* (YcsF, 2P58_B, probability 98.24%), *Salmonella* (PrgI, 2X9C_A, probability 98.19%), *Shigella* (MxiH, 6ZNI_C, probability 98.19%) and *Burkholderia* (BsaL, 2G0U_A, probability 97.24%). We therefore be used Vpa1343 as an indicator of VtrB-mediated production and assembly of the T3SS2 apparatus. We chose an established thiol chemistry-based approach to specifically tag and visualize the formation of T3SS2 needles (*18*). *V. parahaemolyticus* strains expressing Vpa1343 needle proteins with serine to cysteine mutations (S3C, S32C, S62C, S85C and S91C) were tested for the functionality of their T3SS2 apparatuses in terms of effector secretion (Fig. 3A) and relative cytotoxicity in HeLa cells (Fig. 3B). Collectively, these data indicate that among the tested mutants, Vpa1343-S3C was able to produce functional T3SS2 needle filaments and could therefore be tagged with fluorophore-conjugated, thiol-reactive dyes for confocal imaging. The *parSMT1*-ParBMT1 locus-tagging and VtrB-msfGFP constructs were reconstituted in the *V. parahaemolyticus* CAB2 Δ*vpa1343::vpa1343-S3Ck* background, and cells of the resultant strains were grown in either inducing (100 µM TDC) or non-inducing conditions. In both these strains, the Alexa Fluor™ 647 C_2_ Maleimide-stained T3SS2 needles (cyan) were observed to form on the outer membrane near the inner membrane localization of both the *vtrB* genomic locus (Fig. 3C, green) and the VtrB protein (Fig. 3D, blue) in inducing conditions. These needle filaments, however, were not observed in the absence of bile acid induction (Fig. 3C and D). These results demonstrate the second transertion step where the membrane-localized master regulator of T3SS2, VtrB, initiates the synthesis and assembly of the T3SS2 apparatus in its immediate vicinity.

**Figure 3.**
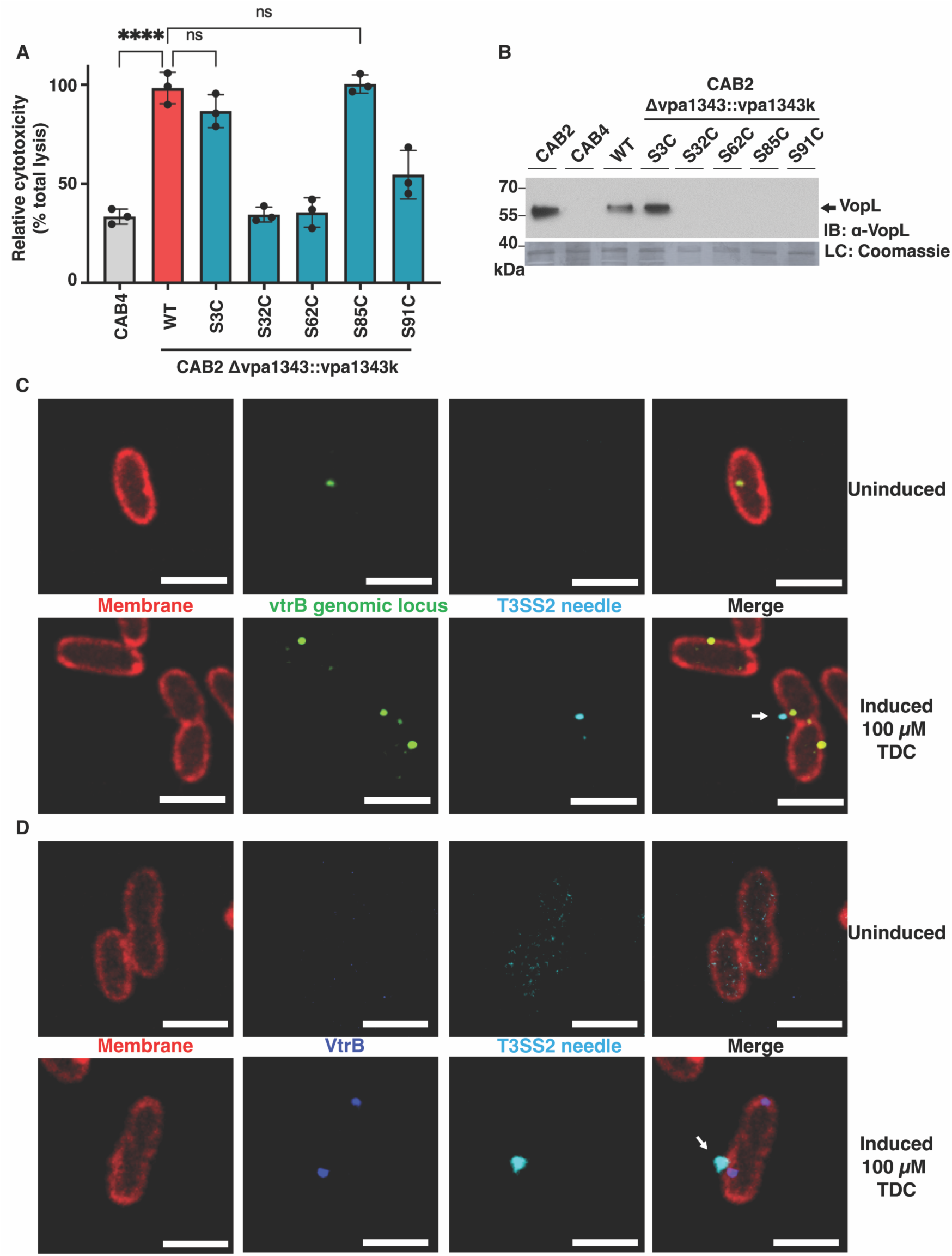
The T3SS2 apparatus, VtrB and *vtrB* genomic locus cluster at the membrane. **A**. Bar graph depicting relative cytotoxicity of *V. parahaemolyticus* CAB2 (WT), CAB4 (T3SS2^-^) and T3SS2 needle mutants (*vpa1343S3C, vpa1343S32C, vpa1343S62C, vpa1343S85C*, and *vpa1343S91C*) in Hela cells. The data represents experiments performed in triplicates with at least two biological replicates (**** represents p < 0.0001, ns = not significant) **B**. Immunoblot for the T2SS2-associated VopL effector, secreted by the strains mentioned above under inducing conditions (0.05% bile). Blot was stained with α-VopL (1:1000) rabbit primary antibodies and α-rabbit HRP-conjugated (1:10,000) secondaries. Identically loaded, Coomassie-stained gel was used as the loading control. **C**. Confocal micrographs of *V. parahaemolyticus* VPKK6 cells cultured in inducing (100 µM TDC) and non-inducing conditions, showing colocalization of T3SS2 needles (false-colored cyan) with the membrane captured *vtrB* locus (green) at the membrane (red). **D**. Confocal micrographs of *V. parahaemolyticus* SC288 cells cultured in the same conditions mentioned above, displaying colocalization of VtrB (false-colored blue) with T3SS2 needles (false-colored cyan) at the membrane (red). **C and D**. White arrows point to T3SS2 needles. Scale bar = 2 µm.

T3SSs are large molecular machines that are well conserved amongst several enteric Gram-negative bacteria and the sheer complexity of these structures warrants tight regulation of their expression and assembly to correspond to host-specific cues (*19*). Bacteria have evolved sophisticated signaling mechanisms utilizing one- and two-component systems to serve this regulatory purpose, thus allowing the bacteria to sense and adapt to everchanging host conditions (*1, 20*). While subsets of the transertion phenomenon have been previously explored by multiple studies (*8, 21-24*), here we have provided the first known direct experimental evidence, demonstrating transertion’s role in coupling membrane-bound, one-component signal transduction cascades with the localized deployment of a large virulence system in response to an external stimulus. In addition to lending credence to the notion that membrane-bound transcription factors exist for precise spatiotemporal regulation of bacterial molecular machines, these findings set the stage for a novel area of study exploring biosynthetic components used to assemble molecules for production of membrane bound complexes.

## Acknowledgments

We thank the Orth lab members for discussions and editing.

## Funding

Welch Foundation grant I-1561

Once Upon a Time…Foundation

National Institutes of Health Grants R35 GM130305 (to K.O) and R35GM128674 (to A.B.D)

The content is solely the responsibility of the authors and does not necessarily represent the official views of the NIH.

K.O. is a W.W. Caruth, Jr. Biomedical Scholar with an Earl A. Forsythe Chair in Biomedical Science.

## Author contributions

K.G.K., S.C., S.S., and K.O. designed the experiments; K.G.K. S.C. S.S. and N.G.R. conducted experiments; K.G.K. performed microscopy; K.G.K. S.C. S.S. and N.G.R. performed cloning and strain preparation; K.G.K and K.O. wrote the manuscript with input from all authors

## Data and materials availability

All materials developed in this study will be made available upon request.

## Supplementary Materials

Materials and Methods

Figs. S1 and S2

Tables S1

References (5, 11, 25-27)

