## Supplementary material for "Transertion is used for localized expression and assembly of *Vibrio parahaemolyticus* T3SS2": Supp File

**Transertion: Localization, Expression and Assembly of the *Vibrio parahaemolyticus* T3SS2**

**This PDF file includes:**

Materials and Methods

Figs. S1 and S2

Tables S1

Materials and Methods

Bacterial strains and media

All bacterial strains used in this study are listed in Table S1. Media was procured from Fisher Scientific™ and chemical reagents from Sigma-Aldrich, unless mentioned otherwise. *E. coli* and *V. cholerae* strains were cultured at 37°C in Miller’s Luria Bertani (LB) broth with agitation. When necessary, the media was supplemented with either 50 µg/ml kanamycin, 25 µg/ml chloramphenicol or 50 µg/ml zeocin. *V. parahaemolyticus* strains were routinely cultured in LB broth supplemented with 3% (w/v) NaCl (MLB) at 30°C, unless noted differently, and when required were supplemented with either 250 µg/ml kanamycin or 25 µg/ml chloramphenicol.

Plasmid Construction

For extrachromosomal expression of gene fragments, the pLAU53 vector backbone was used. Firstly, the vector was PCR linearized with primers Lf/pLAU53_nobla_KpnI and Ri/pLAU53_nobla_SpeI to remove the Amp^R^ cassette. Then, the Kan^R^ cassette was PCR amplified from the pET28a vector (F/pET28a_KmR_KpnI & R/pET28a_KmR_SpeI), restriction digested with KpnI/SpeI and ligated into the linearized vector that had been similarly digested to give pLAU(Kan). SS2A5 was constructed by amplifying the *ygfp-parBMT1* gene fragment from TND1379 chromosomal DNA using primers F/ygfp-parBMT1_NheI and R/parBMT1_HindIII, NheI/HindIII digesting both the fragment as well as the pLAU(Kan) vector, and ligating the resultant fragment into the linearized vector backbone. For SS1I9, the synthesized 1.42 Mbp-*parSMT1* gene fragment was digested out of SS1I3 using enzymes SacI/SalI and ligated into the linearized pDM4 backbone (SacI/SalI). To clone the 0.458 Mbp-*parSP1* fragment into pDM4, primers KK23/KK45 and KK28/KK46 were first used to PCR amplify approximately 1 Kbp regions flanking the chromosomal site of insertion and included the *parSP1* sequence. These two fragments were then integrated into the PCR linearized pDM4 backbone (KK21/KK22) using Gibson Assembly to make SC300. ECKK7 [*cfp-parBP1-ygfp-parBMT1* from TND1379 (KK15/KK16) into pLAU(Kan) (KK13/KK14)], and SC299 [approximately +/- 1 Kbp *vpa1348* from CAB2 (KK52/KK57 and KK55/KK56) and *msgfp* from pC014-*lwcas13a-msfgfp* (KK53/KK54) into pDM4 (KK49/KK58)] were constructed using a similar Gibson Assembly-based strategy. To generate the constructs for Cys-labeling Vpa1343, *vpa1343* including 3 Kbp upstream and downstream flanking regions were first amplified using primers F/pUC18-1343_BamHI and R/pUC18-1343_EcoRI, digested with BamHI/EcoRI and ligated into the pUC18 backbone. To aid in PCR screening of subsequent needle mutants, a KpnI restriction site was introduced into *vpa1343* at T153 using site-directed mutagenesis (F/sdm1343_KpnI and R/sdm1343_KpnI). The plasmid was digested with BglII and the resultant 1.3 Kbp *vpa1343-kpnI* was cloned into the pDM4 backbone (SS1G7). Successive Vpa1343 Cys-labeling constructs were made using site-directed mutagenesis with their corresponding primers detailed in Table S1.

Strain construction and allelic exchange

Both pLAU(Kan) and pDM4-based vectors were inserted into their corresponding recipient *V. parahaemolyticus* strains via triparental conjugation facilitated by *E. coli* DH5α (pRK2043). The transconjugants were selected on minimal marine medium (MMM) agar plates containing either 250 µg/ml kanamycin or 25 µg/ml chloramphenicol for pLAU(Kan) or pDM4-derived constructs, respectively, and confirmed by PCR. Allelic exchange for in-frame deletions/additions/substitutions in *V. parahaemolyticus* strains using pDM4-based constructs were carried out by subsequent growth on MMM agar plates supplemented with 15% (w/v) sucrose for counterselection. The resultant mutants were verified by PCR and sequencing.

T3SS2 expression and effector secretion assay

*V. parahaemolyticus* strains were grown overnight in MLB at 30°C and were diluted the following day to OD_600_ = 0.3 in fresh media. T3SS2 expression was induced by supplementing the media with 0.05% bile salts and incubating the cultures for 3 h at 37°C (*5*). Equal volumes of bacterial cultures, normalized to an OD_600_ = 0.5 were centrifuged at 4000 × g for 10 mins. The pellet/expression fraction was resuspended in 2x Laemmli buffer. The supernatant/secretion fractions were filtered through a 0.22 µM filter and precipitated using 150 µg/ml deoxycholate and overnight treatment with 7.5% (v/v) trichloroacetic acid at 4°C. The following day, the precipitated proteins were collected by centrifuging at 16000 × g for 15 mins, washing the pellets twice in acetone and resuspending them in 2x Laemmli buffer. Western blot analysis was used to detect expression and secretion levels.

Lactate dehydrogenase (LDH) cytotoxicity assay

HeLa cells were plated at 7 × 10^4^ cells per well in a 24-well tissue culture plate in triplicate and were allowed to grow for 16-18 h. Bacterial cultures were induced for T3SS2 expression as described previously by growing the cells for 90 min at 37°C. The induced *V. parahaemolyticus* cultures were added to the HeLa cells at an MOI of 10 and the infection was synchronized by immediately spinning down the plate at 1000 × g for 5 min. 100 µg/ml gentamycin was added to the HeLa cells 2 h post-infection. 200 µl of the spent media from each well was collected 7 h after gentamycin treatment and centrifuged in a 96-well plate at 1000 × g for 5 min. 100 µl of the resultant supernatant was assayed for host cell lysis using the cytotoxicity detection kit (Takara Bio) as per the instructions of the manufacturer. As a positive control, HeLa cells from the uninfected control well were treated with 1% Triton X-100 for 10 min at the end of the gentamycin treatment. HeLa cell lysis by the *V. parahaemolyticus* strains was expressed as a % of cell lysis induced by Triton treatment.

Fluorescence labeling and super-resolution microscopy

T3SS2 expression in *V. parahaemolyticus* strains was initiated in a manner similar to that mentioned above, using 100 µM taurodeoxycholate (TDC) as the inducing bile salt and incubating at 37°C for 25 min. When needed, the media was additionally supplemented with 0.02% arabinose to express the ParB fusion protein of the genomic locus-tagging system. Nile Red (1 µg/ml) and Alexa Fluor™ 647 C_2_ Maleimide (10 µM) were added to the cultures and incubated for a further 20 mins at 37°C to stain the membrane and T3SS2 needles, respectively. The bacterial cells were collected by centrifugation at 6000 × g and fixed at room temperature for 10 min with 3.2% paraformaldehyde. The cell pellets were washed thrice with 1X PBS and were either imaged immediately or stored at 4°C for imaging the following day. 0.5 µl of the cells were spotted on 1% agarose pads mounted onto glass slides and were allowed to dry prior to adding the coverslip. All images were acquired as z-stacks using the Olympus Spin-SR Spinning Disk Confocal Microscope System at 320X magnification (100X oil objective coupled with the 3.2X super-resolution module).

Image processing and data acquisition

All images were processed using the accompanying cellSens Dimension software. Constrained iterative deconvolution (cellSens TruSight 3D Deconvolution module) optimized for super-resolution microscopy (20 iterations) was performed to clean up image noise. The resultant images were used to calculate linear fluorescence intensity profiles (cellSens Count & Measure module).

Statistical Analysis

All data are presented as their mean values ± standard deviation from three or more independent experiments, each conducted in triplicate, unless mentioned otherwise. Unpaired, two-tailed Welch’s t test and one-way ANOVA with Tukey’s multiple comparison tests were used to perform statistical analyses, with a p value < 0.05 being considered as significant.

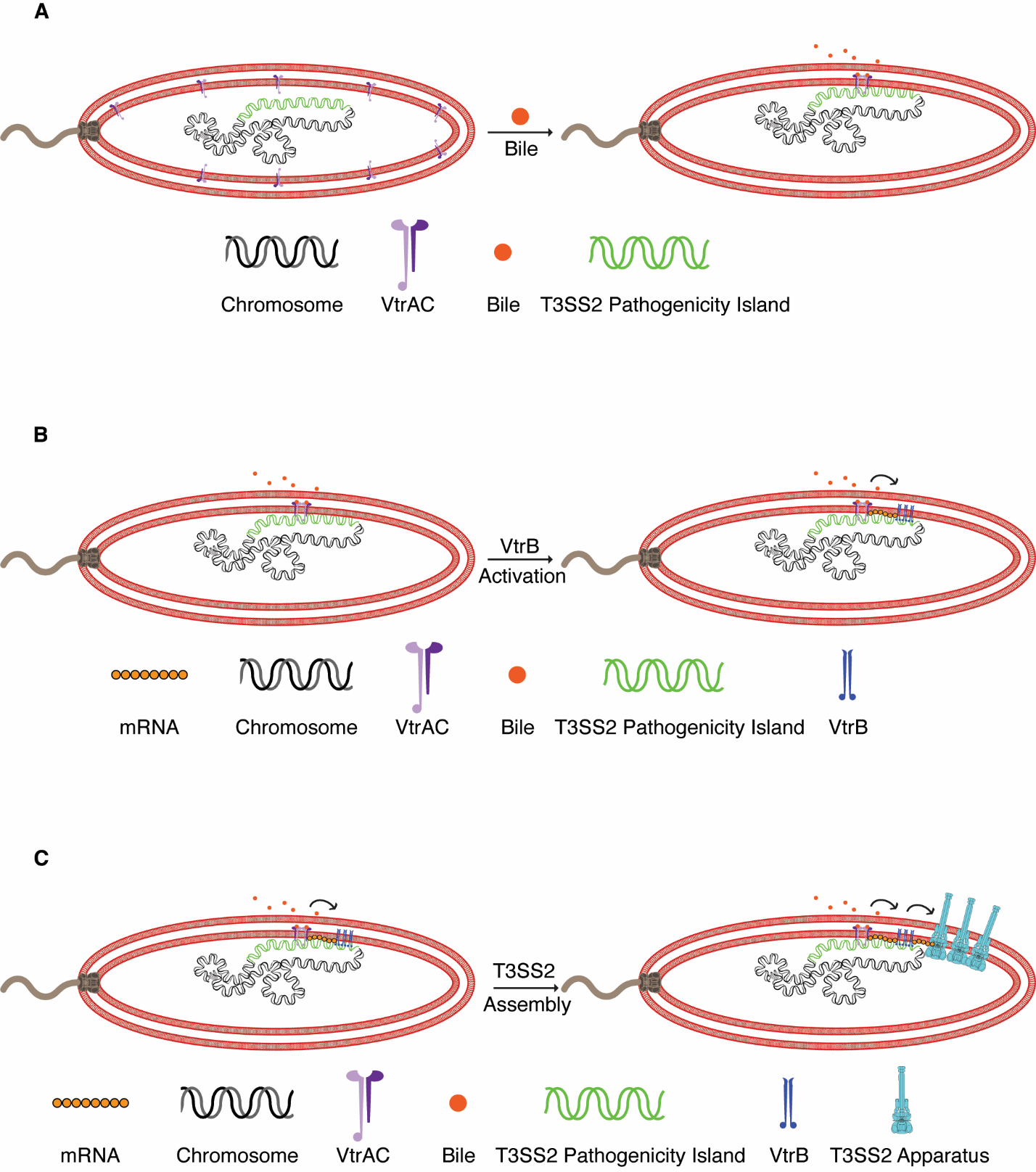

Figure S1. Two-step transertion model for the localized membrane assembly of the T3SS2 apparatus in *V. parahaemolyticus*

Illustrations depicting **A.** Dimerization and activation of VtrA/VtrC upon binding bile acid, which captures the T3SS2 pathogenicity island containing the *vtrB* genomic locus at the inner membrane, **B.** First transertion step, starting with transcription initiation of *vtrB* by VtrA/VtrC, and concurrent translation and membrane insertion of the VtrB protein in the immediate vicinity of the membrane-captured *vtrB* genomic locus, **C.** Second transertion step, starting with transcription initiation of T3SS2 structural and chaperone-associated effector genes by clustered VtrB upon binding to their respective membrane-proximal promoters, and simultaneous translation and membrane assembly of T3SS2 apparatus.

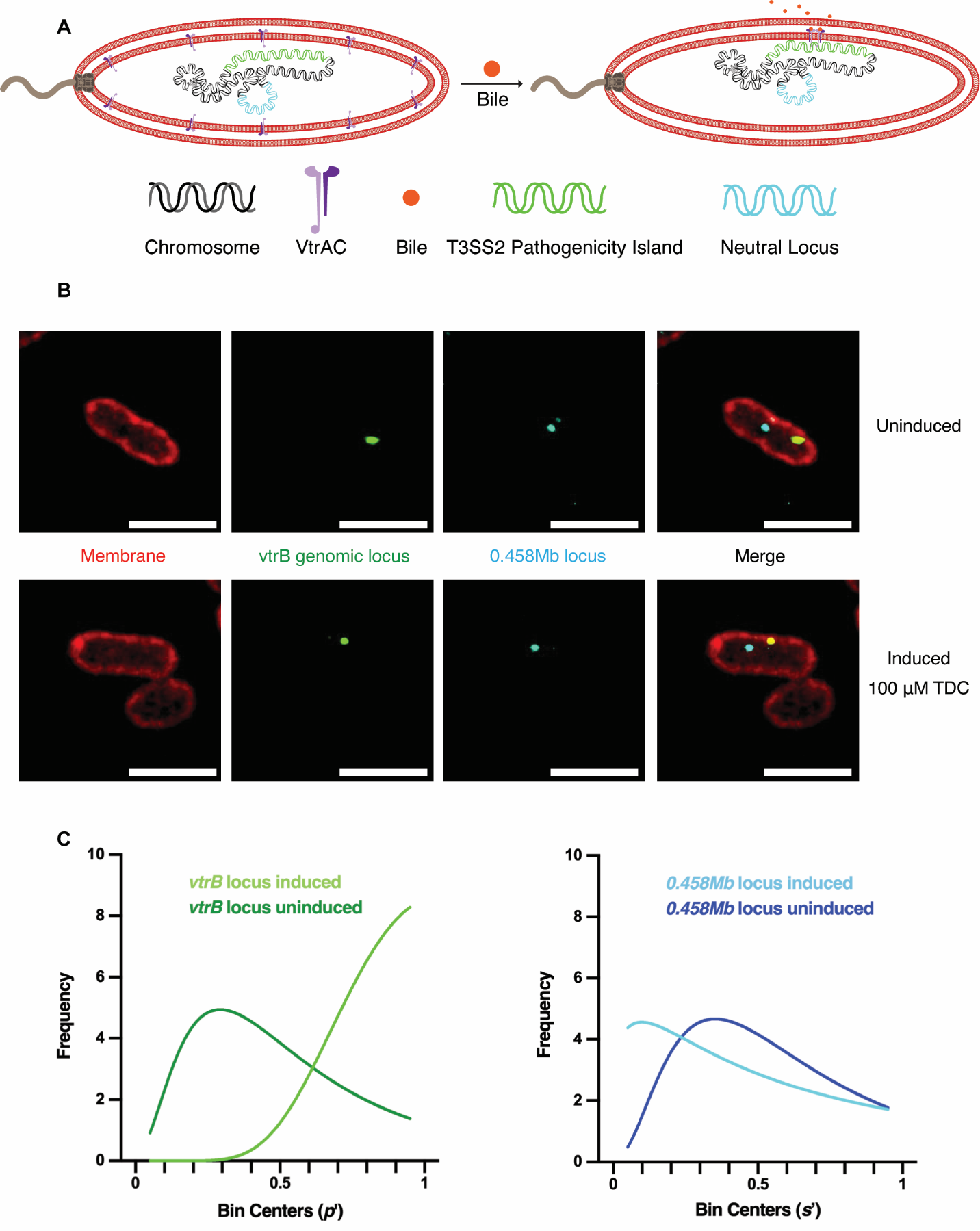
**Figure S2. Bile-induced genomic target capture by activated VtrA/VtrC at the membrane is specific to the *vtrB* locus.**

**A.** Illustration depicting specific capture of the *vtrB* genomic locus, along with the T3SS2 pathogenicity island, at the membrane by bile-activated dimeric VtrA/VtrC, without similarly recruiting a neutral locus lying outside this region. **B.** Confocal micrographs of *V. parahaemolyticus* VPKK11 cells cultured in either non-inducing or inducing (100 µM TDC) conditions, displaying localization of the *vtrB* (green) and neutral (cyan) loci relative to the membrane (red). Scale bar = 2 µm. **C.** Frequency distribution plots with nonlinear regression (lognormal) analyses of normalized *vtrB* loci (p’) and neutral loci (s’) distances in *V. parahaemolyticus* VPKK11 cells, cultured as described above. N=30/locus/condition.

Table S1. Bacterial strains, plasmids and primers.

|  | Relevant characteristics | Source/  Reference |
| --- | --- | --- |
| Strains |  |  |
| *V. parahaemolyticus* |  |  |
| CAB2 | Reference strain, RIMD2210633 Δ*tdh* Δ*trh* Δ*exsA* | (*25*) |
| CAB4 | RIMD2210633 Δ*tdh* Δ*trh* Δ*exsA* Δ*vtrA* | (*25*) |
| SS2B1 | CAB2, P*_araBAD_*-*ygfp-parBMT1*, Km^R^ | This work |
| SS2B3 | CAB2 1.42 Mbp*::parSMT1* | This work |
| SS2B5 | CAB2 1.42 Mbp*::parSMT1*, P*_araBAD_*-*ygfp-parBMT1*, Km^R^ | This work |
| SS1H4 | CAB2 *Δvpa1343::vpa1343-kpnI* | This work |
| SS2B8 | CAB2 *Δvpa1343::vpa1343S3C-kpnI* | This work |
| SS1G6 | CAB2 *Δvpa1343::vpa1343S32C-kpnI* | This work |
| SS1I6 | CAB2 *Δvpa1343::vpa1343S62C-kpnI* | This work |
| SS1H5 | CAB2 *Δvpa1343::vpa1343S85C-kpnI* | This work |
| SS1I7 | CAB2 *Δvpa1343::vpa1343S91C-kpnI* | This work |
| SS2D4 | CAB2 *Δvpa1343::vpa1343S3C-kpnI,* 1.42 Mbp*::parSMT1* | This work |
| VPKK6 | CAB2 *Δvpa1343::vpa1343S3C-kpnI,* 1.42 Mbp*::parSMT1*, P*_araBAD_*-*ygfp-parBMT1*, Km^R^ | This work |
| SC279 | CAB2 *Δvpa1343::vpa1343S3C-kpnI,* 1.42 Mbp*::parSMT1,* 0.458 Mbp*::parSP1* | This work |
| VPKK11 | CAB2 *Δvpa1343::vpa1343S3C-kpnI,* 1.42 Mbp*::parSMT1,* 0.458 Mbp*::parSP1*, P*_araBAD_*-*cfp-parBP1-ygfp-parBMT1*, Km^R^ | This work |
| SC288 | CAB2 *Δvpa1343::vpa1343S3C-kpnI,* 1.42 Mbp*::parSMT1, Δvpa1348::vpa1348-msfgfp* | This work |
| SC295 | CAB2 *Δvpa1343::vpa1343S3C-kpnI,* 1.42 Mbp*::parSMT1, Δvpa1348::vpa1348-msfgfp*, P*_araBAD_*-*cfp-parBMT1*, Km^R^ | This work |
| *V. cholerae* |  |  |
| TND1379 | 0.11 Mbp*::parSP1,* 1.963 Mbp*::parSMT1,* ∆*lacZ::*P*_lac_-CFP-parBP1 yGFP-parBMT1,* Zeo^R^ | (*11*) |
| *E. coli* |  |  |
| DH5α (pRK2073) | Strain carrying the conjugation helper plasmid, pRK2073 | Lab stock |
| Mach1 | Host for cloning vectors | Lab stock |
| S17-1 | Host for pDM4-based plasmids, *λ pir^+^* | Lab stock |
| Plasmids |  |  |
| pUC18 | Cloning vector, Amp^R^ | Lab stock |
| pDM4 | Suicide vector, *γ ori R6K*, *sacB*, Cm^R^ | Lab stock |
| pC014-*lwcas13a-msfgfp* | msfGFP, Amp^R^ | (*26*) |
| pLAU53 | P*_araBAD_*, Amp^R^ | (*27*) |
| pLAU(Kan) | Amp^R^ cassette replaced with Km^R^ cassette of pET28a, Km^R^ | This work |
| SS2A5 | pLAU(Kan) [P*_araBAD_*-*ygfp-parBMT1*], Km^R^ | This work |
| SS1I3 | CAB2 1.42 Mbp-*parSMT1* cloned into pTWIST by gene synthesis | This work |
| SS1I9 | pDM4 [1.42 Mbp-*parSMT1*], Cm^R^ | This work |
| SC300 | pDM4 [0.458 Mbp-*parSP1*], Cm^R^ | This work |
| ECKK7 | pLAU(Kan) [P*_araBAD_*-*cfp-parBP1-ygfp-parBMT1*], Km^R^ | This work |
| SC299 | pDM4 [+/- 1 Kbp *vpa1348-msfgfp*], Cm^R^ | This work |
| SS1F6 | pUC18 [+/- 3 Kbp *vpa1343*], Amp^R^ | This work |
| SS1F7 | pUC18 [+/- 3 Kbp *vpa1343-kpnI*], Amp^R^ | This work |
| SS1G7 | pDM4 [+/- 1.3 Kbp *vpa1343-kpnI*], Cm^R^ | This work |
| SS2A1 | pDM4 [+/- 1.3 Kbp *vpa1343S3C-kpnI*], Cm^R^ | This work |
| SS1G1 | pDM4 [+/- 1.3 Kbp *vpa1343S32C-kpnI*], Cm^R^ | This work |
| SS2H7 | pDM4 [+/- 1.3 Kbp *vpa1343S62C-kpnI*], Cm^R^ | This work |
| SS1G8 | pDM4 [+/- 1.3 Kbp *vpa1343S85C-kpnI*], Cm^R^ | This work |
| SS2H8 | pDM4 [+/- 1.3 Kbp *vpa1343S91C-kpnI*], Cm^R^ | This work |
| Primers |  |  |
| F/pET28a_KmR_KpnI | 5’-AAAAGGTACCATCCTTTGATCTTTTCTACGGGGTCT-3’ | This work |
| R/pET28a_KmR_SpeI | 5’-AAAAACTAGTGAATTAATTCTTAGAAAAACTCATCGAGCATC-3’ | This work |
| Lf/pLAU53_nobla_KpnI | 5’-AAAAGGTACCAGAGTTTGTAGAAACGCAAAAAGGC-3’ | This work |
| Ri/pLAU53_nobla_SpeI | 5’-AAAAACTAGTCTGTCAGACCAAGTTTACTCATATATACTTTAG-3’ | This work |
| F/ygfp-parBMT1_NheI | 5’- AAAAGCTAGCAGGAGGAATTCACC-3’ | This work |
| R/parBMT1_HindIII | 5’-AAAAAAGCTTTTACTCACCTGATTCTGGAAGTC-3’ | This work |
| KK13 | 5’-TTCCAGAATCAGGTGAGTAAAAGCTTGGCTGTTTTGGCGGA-3’ | This work |
| KK14 | 5’-AGTTCTTCTCCTTTACTCATGTGAATTCCTCCTGCTAGAGAGCT-3’ | This work |
| KK15 | 5’-CTCTAGCAGGAGGAATTCACATGAGTAAAGGAGAAGAACTTTTCACTGG-3’ | This work |
| KK16 | 5’-CCGCCAAAACAGCCAAGCTTTTACTCACCTGATTCTGGAAGTCTTTCC-3’ | This work |
| KK21 | 5’-CCGGGTACCATTTTATTTCTGGCGTGGGCTAGTTGTTGATTGG-3’ | This work |
| KK22 | 5’-CGAAAATCAAGCTTAGCATGCATGGCTATAATAGTACTTGAGAAGGAGGCA-3’ | This work |
| KK23 | 5’-CTAGTCTAGATTATGACGTGTATTTCATTATCGATTTTAAATCAAGAACAAG-3’ | This work |
| KK28 | 5’-CATGCCATGGCTATAATAGTACTTGAGAAGGAGGCA-3’ | This work |
| KK45 | 5’-GGTTCAACAAAATAATCGACGCGTCTGCAGAAG-3’ | This work |
| KK46 | 5’-AAAGAAGGATGTTTTCGTCATATGGATCCGATATCGCCG-3’ | This work |
| KK49 | 5’-CTGCAGACGCGTCGATTATTTTGTTGAACCTTTTTGATCAATTCCATAAGGTAAAAC-3’ | This work |
| KK52 | 5’-CGGATCCATATGACGAAAACATCCTTCTTTCTTTTATTCTGAAGTGAATCAC-3’ | This work |
| KK53 | 5’-ATTAGGCGCATTTCTGAACCGACTTCTCCTTTTTCG-3’ | This work |
| KK54 | 5’-GAACCGCAATAAGAAGGATATGGATCTGGAGCTGTAATATAAAAAC-3’ | This work |
| KK55 | 5’-TATCCTTCTTATTGCGGTTCCTACAACTAGTGGAACT-3’ | This work |
| KK56 | 5’-CCCACATTATCTGTCAATCGCTTATAAGATAGAATTTAAAATTTTTGTCACTTCTGC-3’ | This work |
| KK57 | 5’-CGATTGACAGATAATGTGGGTAGCACAACGGCGT-3’ | This work |
| KK58 | 5’-GGTTCAGAAATGCGCCTAATTTTTTAGCTTTGTCCG-3’ | This work |
| F/pUC18-1343_BamHI | 5’-AAAAAGGATCCGACTATGTCGATATGAGTAGTTTTGTAAAAGCTGTC-3’ | This work |
| R/pUC18-1343_EcoRI | 5’-AAAAAGAATTCCCTCATTATCATCTTCATCTAGAGACTCCTCAAC-3’ | This work |
| F/sdm1343_KpnI | 5’-AGCTCCAATCGACCATTGGTACCTGCTTTGTCATG-3’ | This work |
| R/sdm1343_KpnI | 5’-CATGACAAAGCAGGTACCAATGGTCGATTGGAGCT-3’ | This work |
| F/sdm1343_S3C | 5'-ACCTGCACCAGCGTTACATAACATATAAAAATCTCCTATAGCA-3' | This work |
| R/sdm1343_S3C | 5'-TGCTATAGGAGATTTTTATATGTTATGTAACGCTGGTGCAGGT-3' | This work |
| F/sdm1343_S32C | 5'-CTTATCAGCTTCTTTAATCAAACATTCGAAAGAAACACCAACCTCTG-3' | This work |
| R/sdm1343_S32C | 5'-CAGAGGTTGGTGTTTCTTTCGAATGTTTGATTAAAGAAGCTGATAAG-3' | This work |
| F/sdm1343_S62C | 5'-CTTTCTGCCGGGGAATGTTTGCAGCTCCAAC-3' | This work |
| R/sdm1343_S62C | 5'-GTTGGAGCTGCAAACATTCCCCGGCAGAAAG-3' | This work |
| F/sdm1343_S85C | 5'-AACTGGTACTGCTACGTTGAAGTGCATTAAAGATTCTATCTCTTCAGC-3' | This work |
| R/sdm1343_S85C | 5'-GCTGAAGAGATAGAATCTTTAATGCACTTCAACGTAGCAGTACCAGTT-3' | This work |
| F/sdm1343_S91C | 5'-GTTGAAGTCAATTAAAGATTCTATCTGTTCAGCAGCACGTAACATC-3' | This work |
| R/sdm1343_S91C | 5'-GATGTTACGTGCTGCTGAACAGATAGAATCTTTAATTGACTTCAAC-3' | This work |
